# Parkinson disease age of onset GWAS: defining heritability, genetic loci and a-synuclein mechanisms

**DOI:** 10.1101/424010

**Authors:** Cornelis Blauwendraat, Karl Heilbron, Costanza L. Vallerga, Sara Bandres-Ciga, Rainer von Coelln, Lasse Pihlstrøm, Javier Simón-Sánchez, Claudia Schulte, Manu Sharma, Lynne Krohn, Ari Siitonen, Hirotaka Iwaki, Hampton Leonard, Alastair J. Noyce, Manuela Tan, J. Raphael Gibbs, Dena G. Hernandez, Sonja W. Scholz, Joseph Jankovic, Lisa M. Shulman, Suzanne Lesage, Jean-Christophe Corvol, Alexis Brice, Jacobus J. van Hilten, Johan Marinus, The 23andMe Research Team, Pentti Tienari, Kari Majamaa, Mathias Toft, Donald G. Grosset, Thomas Gasser, Peter Heutink, Joshua M Shulman, Nicolas Wood, John Hardy, Huw R Morris, David A. Hinds, Jacob Gratten, Peter M. Visscher, Ziv Gan-Or, Mike A. Nalls, Andrew B. Singleton, for the International Parkinson’s Disease Genomics Consortium (IPDGC)

## Abstract

Increasing evidence supports an extensive and complex genetic contribution to Parkinson’s disease (PD). Previous genome-wide association studies (GWAS) have shed light on the genetic basis of risk for this disease. However, the genetic determinants of PD age of onset are largely unknown. Here we performed an age of onset GWAS based on 28,568 PD cases. We estimated that the heritability of PD age of onset due to common genetic variation was ~0.11, lower than the overall heritability of risk for PD (~0.27) likely in part because of the subjective nature of this measure. We found two genome-wide significant association signals, one at *SNCA* and the other a protein-coding variant in *TMEM175*, both of which are known PD risk loci and a Bonferroni corrected significant effect at other known PD risk loci, *INPP5F/BAG3*, *FAM47E/SCARB2*, and *MCCC1.* In addition, we identified that *GBA* coding variant carriers had an earlier age of onset compared to non-carriers. Notably, *SNCA*, *TMEM175*, *SCARB2*, *BAG3* and *GBA* have all been shown to either directly influence alpha-synuclein aggregation or are implicated in alpha-synuclein aggregation pathways. Remarkably, other well-established PD risk loci such as *GCH1, MAPT* and *RAB7L1*/NUCKS1 (PARK16) did not show a significant effect on age of onset of PD. While for some loci, this may be a measure of power, this is clearly not the case for the *MAPT* locus; thus genetic variability at this locus influences whether but not when an individual develops disease. We believe this is an important mechanistic and therapeutic distinction. Furthermore, these data support a model in which alpha-synuclein and lysosomal mechanisms impact not only PD risk but also age of disease onset and highlights that therapies that target alpha-synuclein aggregation are more likely to be disease-modifying than therapies targeting other pathways.

## Introduction

Parkinson disease (PD) is the most common neurodegenerative movement disorder. PD is pathologically characterized by the loss of dopaminergic neurons in the substantia nigra and α-synuclein (encoded by *SNCA*) protein aggregates. Current estimates are that in 2015 there were 6.9 million PD patients worldwide and this number is predicted to double to 14.2 million in 2040 (Dorsey and Bloem, 2018). PD has a strong age-dependent prevalence and male-female differences, where males are ~1.5 times more likely to develop PD (Moisan et al., 2016; Reeve et al., 2014).

The exact cause of PD is unknown, however there is clear evidence that genetic variability plays a role in both disease development and progression. In the past two decades, mutations in several genes have been identified that cause monogenic forms of PD, accounting for about 1-5% of all PD cases (Singleton and Hardy, 2016). The majority of PD cases are therefore referred to as sporadic PD. Genome-wide association studies have successfully identified over 40 loci robustly associated with PD. Interestingly, variability in several genes, including *SNCA*, *LRRK2*, *VPS13C*, *GCH1* and *GBA* appear to play a role in both monogenic and sporadic disease (Chang et al., 2017).

Substantial evidence suggests that genetic variation plays a role in age of onset where, for example, monogenic forms of PD often present as early onset cases, but the identity of the exact genetic component in sporadic PD remains unclear (Hamza and Payami, 2010; Nalls et al., 2015). Additionally, there is some evidence that age of onset may be different between males and females (Alves et al., 2009; Haaxma et al., 2007). Multiple studies have nominated variants and genes of interest, but replication and large cohort studies are lacking (Hill-Burns et al., 2016; Latourelle et al., 2009; Nalls et al., 2015). Additionally, rare coding variants in PD risk genes like GBA and LRRK2 have been associated with an earlier age of onset (Gan-Or et al., 2015; Lee et al., 2017; Malek et al., 2018; Marder et al., 2015; Nichols et al., 2009; Xiao et al., 2018). Currently the best predictor of PD age of onset is a genetic risk score (GRS) based on cumulative genetic PD risk (Escott-Price et al., 2015; Nalls et al., 2015), implying broad genetic overlap between PD susceptibility and PD age of onset.

In this study, we performed the largest PD age of onset genome-wide association study (GWAS) to date including 28,568 PD cases. Using this large cohort, we have estimated the heritability of PD age of onset and have identified several associated variants. Additionally, we have shown that the latest PD GRS remains highly correlated with of age of onset.

## Materials and methods

### Processing of IPDGC datasets

Genotyping data (all Illumina platform based) was obtained from International Parkinson’s Disease Genomics Consortium (IPDGC) members, collaborators and public resources (Supplementary Table 1). All datasets underwent quality control separately, both on individual level data and variant level data before imputation. In brief, we excluded individuals with discordance between genetic and reported sex, low call rate (<95%), heterozygosity outliers (F statistic cut-off of > −0.15 and < 0.15), ancestry outliers (+/− 6 standard deviations from means of eigenvectors 1 and 2 of the 1000 Genomes phase 3 CEU and TSI populations from principal components (Genomes Project et al., 2015)), and individuals with cryptic relatedness of > 0.125 using PLINK (version 1.9) (Chang et al., 2015) or GCTA (version 1.02) (Yang et al., 2011) keeping a random proband from any related groups. We further excluded genotypes with a missingness rate of > 5%, a minor allele frequency < 0.001. Remaining samples were submitted per dataset separately to the Michigan imputation server (Das et al., 2016) using Eagle v2.3 imputation (Loh et al., 2016) based on reference panel HRC r1.1 2016 (McCarthy et al., 2016). Principal components (PCs) were created for each dataset from the directly assayed genotypes using PLINK. For the PC calculation, variants were filtered for minor allele frequency (>0.01), genotype missingness (<0.05) and Hardy–Weinberg equilibrium (P =>1E-6). Remaining variants were pruned using a PLINK pairwise pruning (default settings, window size of 50, window shift: 5 SNPs and r2 of 0.5) and PCs were calculated on the pruned variants.

After imputation, all datasets were merged and analyzed for duplicates and cryptic relatedness individuals between datasets (>0.125) using GCTA. One individual was removed at random from pairs of duplicated or related individuals. Due to the large variation in the geographical location / ancestry, genotyping array and average reported age of onset of the IPDGC datasets, we performed genome-wide association analyses for each dataset separately to minimize potential biases and meta-analyzed the results (Supplementary Table 1 and Supplementary Figure 1). Where possible, age of onset was defined based on patient report of initial manifestation of parkinsonian motor signs (tremor, bradykinesia, rigidity or gait impairment). Where this information was not available, age of diagnosis was used as a proxy for onset age. For one study (Oslo Parkinson’s Disease Study) both the age of onset and age of diagnosis was available for a large number of cases (n=321). The Pearson correlation between age of onset and age of diagnosis was 0.965, P<2.2E-16 suggesting that age of diagnosis is a good replacement for age of onset, with an average ~1.97 years delay from age of onset to diagnosis. We excluded individuals with PD onset before the age of 20 or after the age of 100 (see Supplemental Figure 2 of distribution of ages).

The resulting quality controlled and imputed datasets of PD cases (total n=17,415) were analyzed with the formula AGE_AT_ONSET ~ SNP + SEX + PC1-PC5. Analyses were performed per dataset with rvtests linear regression using imputed dosages (Zhan et al., 2016). System Genomics of Parkinson’s Disease (SGPD) data (n=581) was processed as above with the following minor changes: A linear regression model in PLINK was used to run a GWAS for age of onset and relatedness cut-off of 0.05 was used. The association analysis was adjusted for sex and the first 10 eigenvectors from PC analysis.

Additionally, to identify spurious associations due to age-related mortality, we ran GWAS using the controls from the IPDGC datasets (n=16,502). In total 3,083,373 variants passed filter criteria, the genomic inflation factor of the meta-analysis was λ=0.995 (see Supplemental Figure 5 for Manhattan plot and 5 for QQ-plot)

### Processing of the 23andMe dataset

As an independent second cohort we used the 23andMe PD dataset, which consisted of customers of the personal genetics company, 23andMe, Inc., who had consented to participate in research. PD patients were recruited through a targeted email campaign in conjunction with the Michael J. Fox Foundation or other partners, patient groups and clinics. PD cases were individuals who self-reported having been diagnosed with PD and who self-reported their age of PD diagnosis. We used age of diagnosis rather than age of symptom onset because symptom onset is often gradual and may be more difficult to self-report accurately. Similar to the Oslo Parkinson’s Disease Study dataset, the correlation was high between age of diagnosis and age of onset (ρ = 0.917, P<1E-300) and the average difference between age of diagnosis and age of onset was 2.58 years. We excluded individuals if they reported a change in diagnosis or uncertainty about their diagnosis. We have previously shown that self-reported PD case status is accurate with 50 out of 50 cases confirmed via telemedicine interview (Dorsey et al., 2015). To improve data quality and decrease the probability of data entry errors, we excluded individuals who self-reported an age of diagnosis greater than their current age and individuals with an age of diagnosis before the age of 40 (excluded 4.7% of individuals, see Supplemental Figure 3 for the age of diagnosis distribution).

We included unrelated individuals who were genetically inferred to have >97% European ancestry. We did this because 88.9% of all people with PD in the 23andMe database had >97% European ancestry and because PD incidence and prevalence may differ by ethnicity.

We also removed individuals who self-reported having ever been diagnosed with: 1) an atypical parkinsonism or a non-parkinsonian tremor disorder; or 2) stroke, deep vein thrombosis, or pulmonary embolism (to reduce the probability of including individuals with vascular parkinsonism).

DNA extraction and genotyping were performed on saliva samples by Clinical Laboratory Improvement Amendments (CLIA) certified and College of American Pathologists (CAP) accredited clinical laboratories of Laboratory Corporation of America. Quality control, imputation and genome-wide analysis were performed by 23andMe. Samples were genotyped on a 23andMe custom genotyping array platform (Illumina HumanHap550+ Bead chip V1 V2, OmniExpress+ Bead chip V3, custom array V4). Samples had minimum call rates of 98.5%. After quality control, a total of 904,040 SNPs and indels (insertions/deletions) were genotyped across all platforms (for extended genotyping and sample quality details, see (Eriksson et al., 2010; Hyde et al., 2016; Lo et al., 2017). A maximal set of unrelated individuals was chosen for the analysis using a segmental identity-by-descent (IBD) estimation algorithm (Henn et al., 2012) to ensure that only unrelated individuals were included in the sample. Individuals were defined as related if they shared more than 700 cM IBD, including regions where the two individuals shared either one or both genomic segments IBD. This level of relatedness (~20% of the genome) corresponds to approximately the minimal expected sharing between first cousins in an outbred population.

We imputed participant genotype data against the September 2013 release of the 1,000 Genomes phase 1 version 3 reference haplotypes. We phased and imputed data for each genotyping platform separately. We phased using an internally developed phasing tool, Finch, which implements the Beagle haplotype graph-based phasing algorithm (Browning and Browning, 2007), modified to separate the haplotype graph construction and phasing steps. In preparation for imputation, we split phased chromosomes into segments of no more than 10,000 genotyped SNPs, with overlaps of 200 SNPs. We excluded any SNP with Hardy-Weinberg equilibrium P < 10E−20, call rate < 95% or with large allele frequency discrepancies compared to the European 1,000 Genomes reference data. Frequency discrepancies were identified by computing a 2 × 2 table of allele counts for European 1,000 Genomes samples and 2,000 randomly sampled 23andMe customers with European ancestry, and then using the table to identify SNPs with P < 1E−15 by χ2 test. We imputed each phased segment against all-ethnicity 1,000 Genomes haplotypes (excluding monomorphic and singleton sites) using Minimac2 (Browning and Browning, 2007; Fuchsberger et al., 2015), using five rounds and 200 states for parameter estimation. After quality control, which included R2 > 0.5 variant removal, we analyzed 11,956,580 SNPs. Remaining PD cases (n=10,572) were analyzed using the formula AGE_AT_DIAGNOSIS ~ SNP + SEX + PC1-PC5 + genotyping platform. The genomic inflation factor was *λ* = 1.015266.

### Meta-analyzing dataset

The 17 IPDGC datasets were meta-analyzed using METAL (v.2011-03-25) using default settings (Willer et al., 2010). We excluded SNPs with an I^2^ statistic >50% and variants that were present in fewer than 66.7% of the datasets. One genome-wide significant variant was excluded due to this filter criteria (chr4:90711770, I^2^ = 58.1%. This resulted in a total of 6,850,647 variants and a genomic inflation factor of *λ* = 1.001. For the combined GWAS we meta-analysed all 17 IPDGC datasets with the 23andme dataset using the same quality control steps.

### Additional analyses and figures

PD GWAS loci were obtained from (Chang et al., 2017) using Table 1 and Table 2 and using P.META from the provided summary stats (see Supplementary Table 3). Additionally, to assess the influence of a genetic risk score (GRS) for PD case-control status on PD age of onset we obtained the GRS from the most recent PD GWAS Supplementary (Chang et al., 2017). The GRS was calculated and processed using PLINK for each individual as described previously (Nalls et al., 2015). Associations were performed using the formula AGE_AT_ONSET ~ GRS + SEX + PC1-PC5. Quantile-quantile (QQ)-plots and Manhattan plots were generated using FUMA (Watanabe et al., 2017). Forest plots were generated in R using the package rmeta (https://cran.r-project.org/web/packages/rmeta/index.html). Locus plots (Pruim et al., 2010) were generated for the genome-wide significant loci and were compared to the latest published PD GWAS (Chang et al., 2017). GCTA-cojo analyses were performed to identify whether there were multiple independent signals in loci of interest using all IPDGC datasets (excluding SGPD) described in Supplementary Table 1 as a reference panel (Yang et al., 2012; Yang et al., 2011). This reference panel was created using variants passing the following criteria: R^2^<0.8, minor allele frequency > 0.001, Hardy-Weinberg equilibrium *P* < 1E-6, and a maximum variant missingness of 5%. The genetic correlation between the PD age of onset GWAS and the PD GWAS (Chang et al., 2017) was calculated using LD Score Regression (LDSC) (Bulik-Sullivan et al., 2015) using default settings for European populations. Heritability was estimated using LDSC and GCTA based on GWAS summary statistics and individual-level genotypes, respectively.

### Power calculations

We performed power calculations using the method of Brion et al. (Brion et al., 2013)

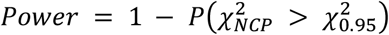

where 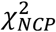 is a random variable from a non-central *χ*^2^ with one degree of freedom, and 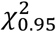 is the threshold of a central *χ*^2^ distribution with one degree of freedom and a type-I error rate of 0.05. The non-centrality parameter was calculated using the method of Sham and Purcell (Sham and Purcell, 2014)

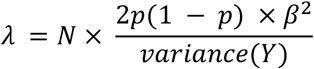

where *λ* is the non-centrality parameter, *p* is the minor allele frequency, *β* is the effect size in years, and variance(*Y*) is the variance in PD age of onset. We estimated variance(*Y*) by taking the mean variance across each cohort, weighted by their sample size. We assessed effect sizes between 0.01 to 1 years and reported the minimum effect size that yielded >80% power.

### Data availability

IPDGC GWAS summary statistics are available (after publication) on the IPDGC website (http://pdgenetics.org/resources) and 23andMe GWAS summary statistics will be made available to qualified researchers under an agreement with 23andMe that protects the privacy of the 23andMe participants. Please visit http://research.23andme.com/collaborate/#publication for more information and to apply to access the data.”

## Results

### Initial data overview

In total we included 18 datasets: the IPDGC data contains of 17 independent cohorts (n=17,996) and the 23andMe PD cohort (n=10,572, see Supplementary Table 1 for more details). The average age of onset in the IPDGC dataset was 62.14 (range 20-96, SD=12.08), while in the 23andMe dataset the average age of diagnosis was 60.71 (range 40-97, SD=9.98). We found minor differences in age of onset in females and males in both the IPDGC dataset (females = 62.15, SD=11.71; males = 62.03, SD=11.95) and the 23andMe dataset (females = 59.95, SD= 9.79; males = 61.20, SD=10.07).

### Heritability of Parkinson’s disease age of onset

Using LDSC, the heritability of PD age of onset was h^2^=0.076 (SE=0.0277) in the IPDGC cohort, h^2^=0.0805 (SE=0.0403) in the 23andMe cohort, and 0.109 (SE=0.0255) in the complete meta-analysis. These heritability estimates for PD age at onset were similar to estimates derived using GCTA in the largest IPDGC dataset (IPDGC NeuroX, h^2^=0.0798, SE=0.0391, P=0.0184, N=5,428) and in the 23andMe dataset (h^2^=0.1235, SE=0.0341, P=1.031E^−4^, N=10,697). Heritability estimates in the other IPDGC datasets were considered less reliable due to the low number of included cases.

### SNCA as top associated loci with Parkinson’s disease age of onset

Two loci reached genome-wide significance, *SNCA* and *TMEM175*, both of which are established PD risk loci based on PD case/control GWASes (Figure 1). Common variation at the *SNCA* locus has clearly been established as a risk factor PD and both rare mutations and whole gene multiplications have been identified to cause monogenic PD (Polymeropoulos et al., 1997; Singleton et al., 2003). Several independent signals have been reported in this locus, where initially a variant in intron 4 was identified in PD (Simón-Sánchez et al., 2009), later a second independent signal at the 5’ end was identified (Nalls et al., 2014), which is also associated with Dementia with Lewy bodies (Guerreiro et al., 2018) and currently at least three independent signals are present in this locus (Pihlstrom et al., 2018). Here, we identified both signals to be associated with age of onset (Figure 2A). The 3’ end signal is the strongest with rs356203 as the most associated SNP, resulting in a reduction of ~0.6 year in age of onset (P_meta=1.90E-12, beta=−0.626, SE=0.0890). Using conditional analysis, we also identified the 5’ end signal (Figure2B, rs983361, 4:90761944, P_conditional=6.82E-6, beta=−0.484, SE=0.108)

**Figure 1:**
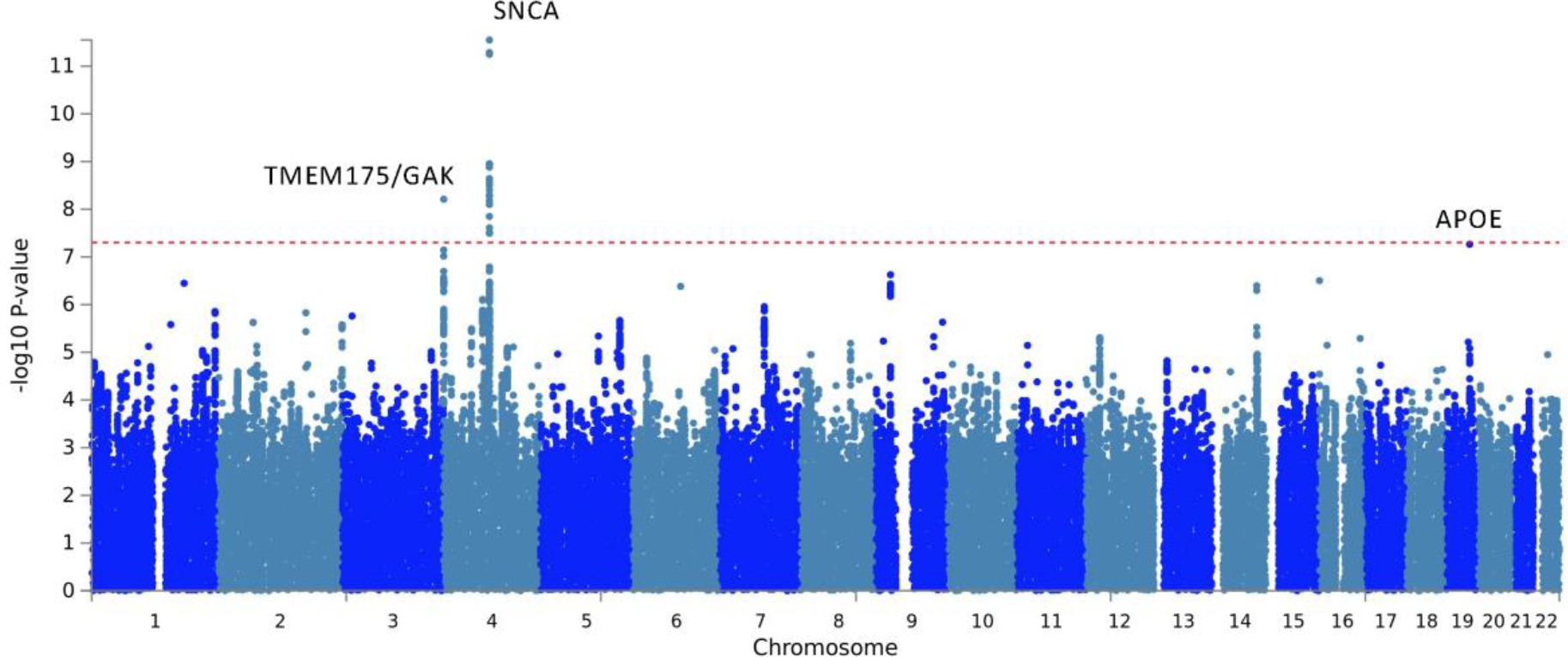
Manhattan plot of Parkinson’s disease age of onset GWASs. Based on meta-analyses of datasets (n=28,568) using 7,426,111 SNPs. Two genome-wide significant loci were identified: *SNCA* and the *TMEM175*/*GAK*. Additionally, one borderline significant locus was found, *APOE*.

**Figure 2:**
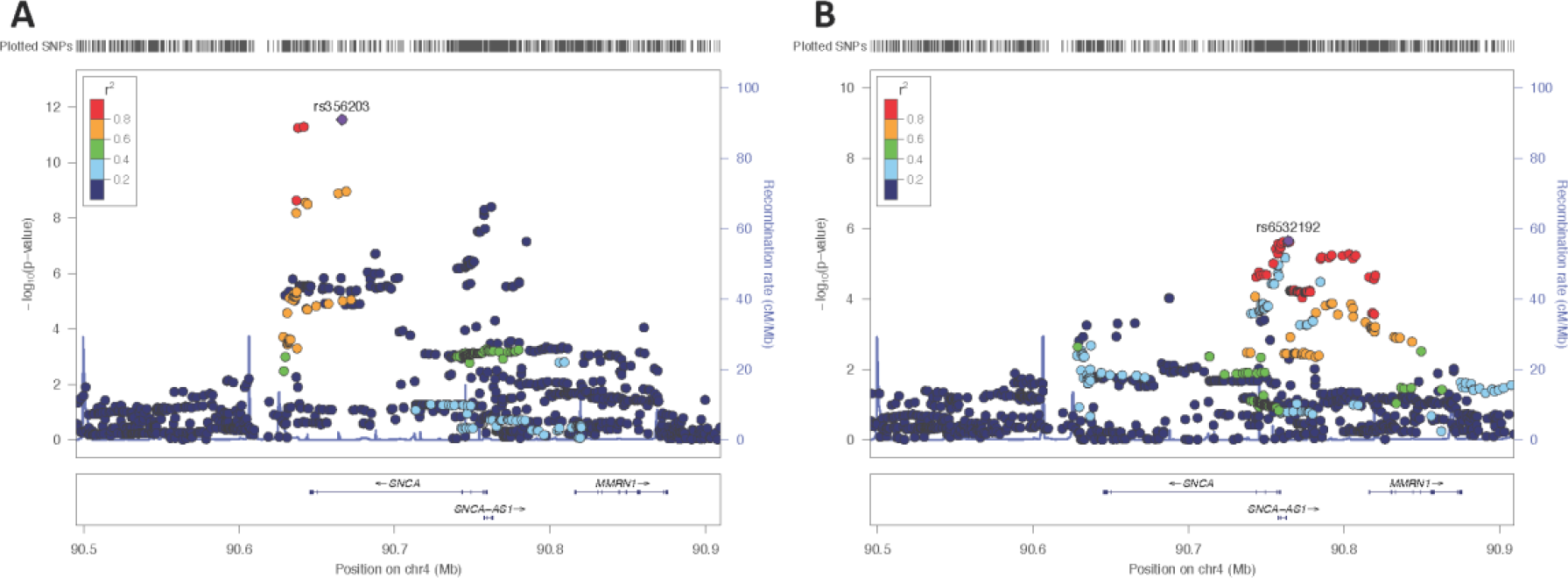
*SNCA* association with Parkinson’s disease age of onset. A) Locus zoom plot of the association signal of the *SNCA* locus. The age of onset association signal is primarily based at the 3’ end of the *SNCA* gene, highly similar as the Parkinson’s disease GWAS signal of Chang et al 2017 (Supplementary Figure 8). B) Conditional analysis based on the most associated SNP (rs356203) shows that there is a secondary signal at the 5’ end of the *SNCA* gene rs6532192, as well similar as previously reported for the Parkinson’s disease GWAS Chang et al 2017.

### A coding variant in TMEM175 is associated with Parkinson’s disease age of onset

The *TMEM175/GAK* locus is another known PD locus where two independent signals have been identified (Nalls et al., 2014). While it is more parsimonious to assume that a single GWAS peak is the product of a single causal gene in the locus, there is evidence that both *TMEM175* and *GAK* may modulate PD risk (Dumitriu et al., 2011; Jinn et al., 2017).*TMEM175* and *GAK* share a promoter, suggesting that they may be co-regulated and may perform related cellular functions. Interestingly, the *TMEM175/GAK* locus is one of the few PD GWAS loci where a coding variant (*TMEM175* p.M393T, exon 11, rs34311866) is amongst the highest associated variants (Figure 2A). This coding variant was also the most associated variant in this locus in our PD age of onset analysis (P_meta=9.62E-9, beta=−0.613, SE=0.107), resulting in an average reduction of ~0.6 years (Figure 3B and Figure 5). Using conditional analysis based on rs34311866 we did not identify a second independent signal in this locus (P=0.879, beta=−0.0240, SE=0.158, Supplementary Figure 9).

**Figure 3:**
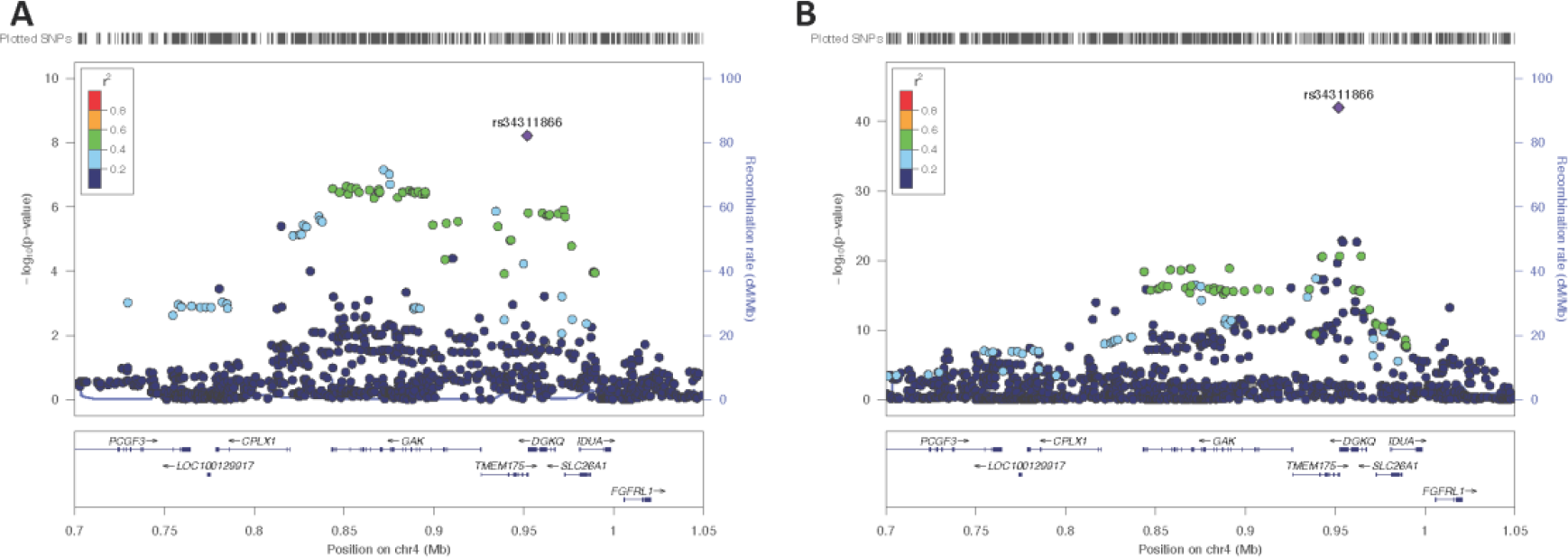
*TMEM175 GAK* locus association with Parkinson’s disease age of onset and Parkinson’s disease. A) Locus zoom plot of the association signal of the *TMEM175*/*GAK* locus with Parkinson’s disease age of onset. The association signal is primarily based on the coding variant *TMEM175* p.M393T, rs34311866. B) Locuszoom plot of the association signal of the *TMEM175*/*GAK* locus with Parkinson’s disease vs controls Chang et al 2017. Similarly, as in A the main signal is based on the coding variant *TMEM175* p.M393T, rs34311866.

**Figure 4:**
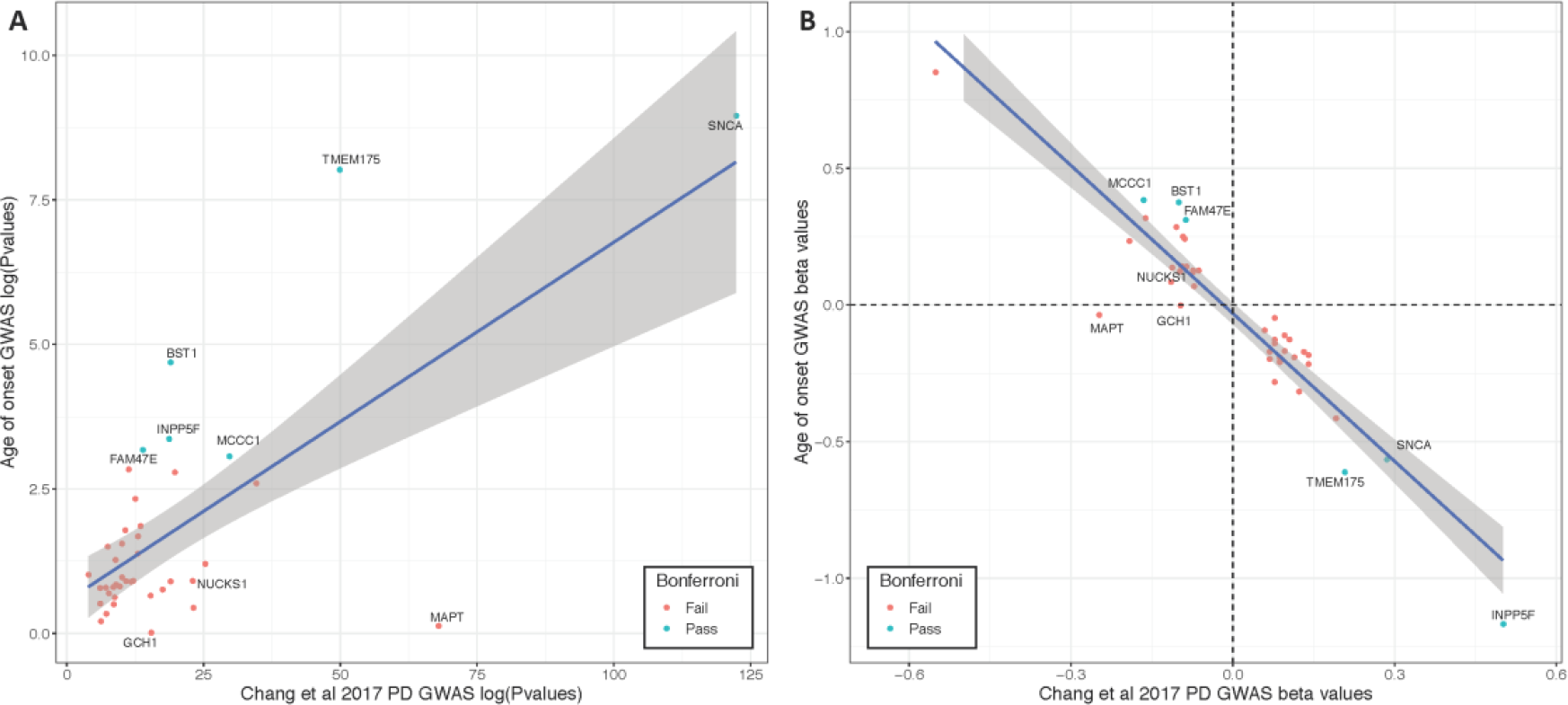
P-value and beta value correlation plot between PD GWAS and age of onset GWAS. A) Log transformed P-values were plotted of the Chang et al 2017 PD GWAS (x-axis) and the current age of onset GWAS (y-axis). SNPs are annotated by their closest gene. Green dots are loci that pass Bonferroni correction and red are loci that did not pass Bonferroni correction. B) Beta-values were plotted of the Chang et al 2017 PD GWAS (x-axis) and the current age of onset GWAS (y-axis). SNPs are annotated by their closest gene. Green dots are loci that pass Bonferroni correction and red are loci that did not pass Bonferroni correction.

**Figure 5:**
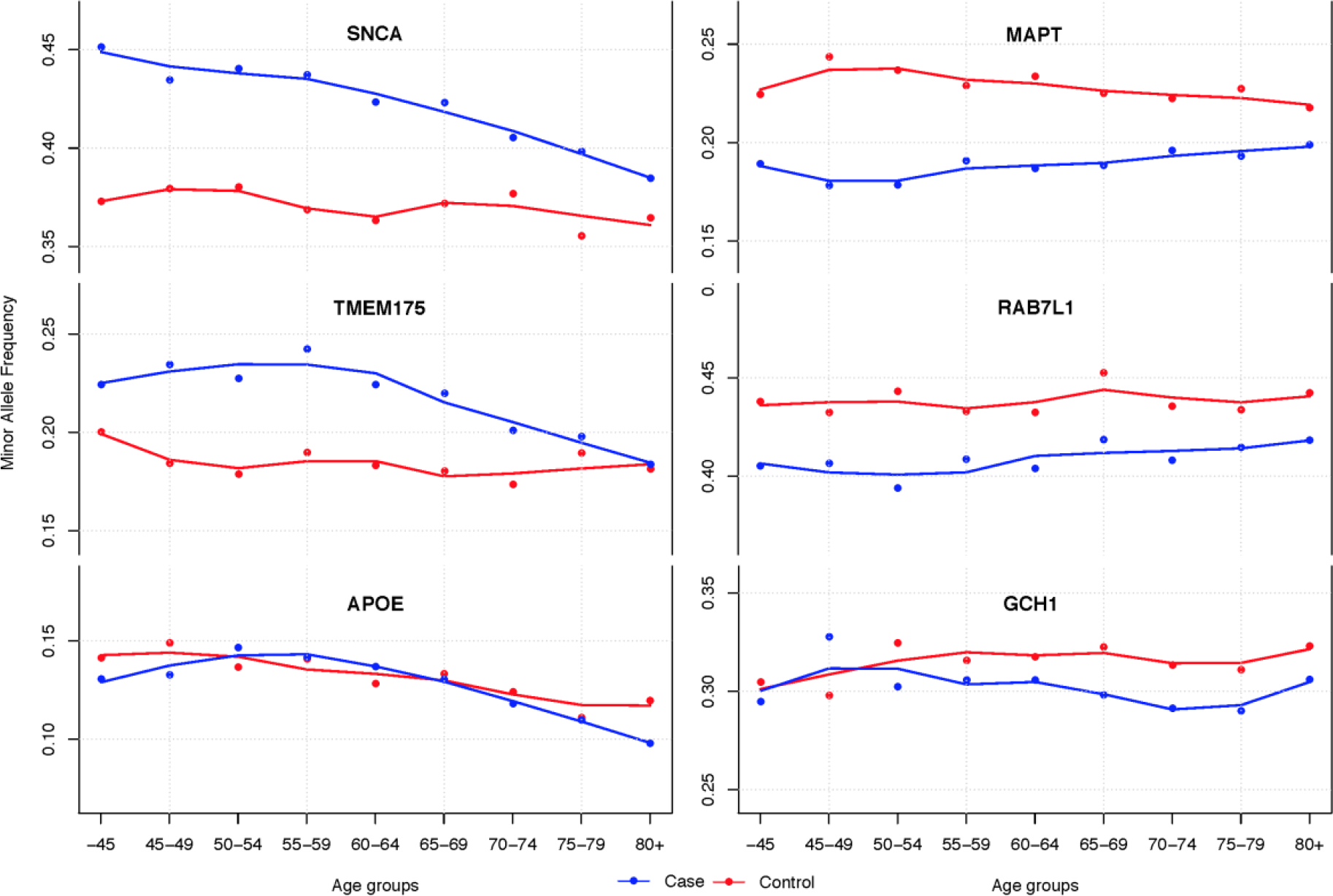
Minor allele frequency differences correlated with age groups separated by case-control status. Genotype data was merged and minor allele frequency were calculated per age group and separated by case control status (IPDGC data only: N_cases=17,996 and N_controls=16,502, each age group contains >850 individuals). Lines were using LOESS regressions based on the minor allele frequency per age group. On the left the three most significant loci associated with PD age of onset are shown; *SNCA*=rs356203, *TMEM175/GAK*=rs34311866 and *APOE*=rs429358. On the right three significant PD case control GWAS loci are shown to have no clear effect on PD age of onset; *MAPT*=rs17649553, *RAB7L1*=rs823118, *GCH1*= rs11158026. Standard error-bars are not shown in figure but are neglectable (~0.015) for each age-group.

### Parkinson’s disease high risk variants in GBA and LRRK2 and age of onset

*LRRK2* p.G2019S and *GBA* variants are the most commonly identified genetic risk factors for PD. *LRRK2* p.G2019S is identified in ~1% of the general European PD population, but higher frequencies have been reported in other populations (Correia Guedes et al., 2010). *GBA* variants are more commonly identified in Ashkenazi Jewish populations but are also present in the European population (Sidransky et al., 2009).

*LRRK2* p.G2019S (rs34637584) has already been described to have large range of age of onset and reduced penetrance (Lee et al., 2017; Marder et al., 2015). Due to the low frequency and low imputation quality of this variant it did not pass the variant level QC and was only tested in 9 of the 18 datasets. Similar to previous studies (Correia Guedes et al., 2010), we identified 140 carriers in the IPDGC datasets (~0.8% of all PD cases). The average age of onset for p.G2019S carriers was 66.58 (range= 36-95, SD=11.66), which is ~4 years later compared to the average age of onset of non-carriers in the IPDGC datasets (62.14, range 20-96, SD=12.08). However, this is based on a relatively small amount of cases and some cohorts excluded *LRRK2* p.G2019S carriers pre-genotyping or excluded familial PD cases. Besides, the non-coding variant at the 5’ end of *LRRK2* (rs76904798) which is also identified as a risk factor PD did not show an association with age of onset (P=0.1267, beta=−0.184, SE=0.121).

For *GBA* there have been several reports that variants in *GBA* affect age of onset in PD (Davis et al., 2016; Gan-Or et al., 2008; Lill et al., 2015). In our large meta-analysis we identified a borderline significant hit for the *GBA* p.N370S variant (rs76763715, *P*=2.628E-6, beta=−2.600, SE=0.553) which resulted in a relatively large reduction of 2.6 year in age of onset. This variant was only tested in 13 of the 18 datasets due to the low frequency or low imputation quality. Using conditional analysis, we did not identify other clear significant signals in the *GBA* gene (Supplementary Figure 11). However, other the coding PD risk variants in *GBA* reached nominally significant *P* values (rs2230288, p.E326K: P_meta=0.00123, beta=−0.929, SE=0.287 and rs75548401, p.T369M: P_meta=0.01153, beta=−1.281, SE=0.507).

### Parkinson’s disease genetic risk loci are associated with age of onset

Previously a GRS based on PD GWAS loci has been identified as the main genetic predictor for age of onset (Nalls et al., 2015). Here we tested the association between the updated 47-SNP GRS used in the latest PD GWAS study (Chang et al., 2017). For each IPDGC dataset the GRS was calculated and we identified a consistent association between the Chang et al. GRS and PD age of onset. After meta-analyzing results from the individual cohorts, we found that each 1SD increase in GRS led to an earlier age of onset by ~0.8 years (summary effect=−0.801, 95%CI (−0.959, −0.643), I2=11.6%) (Figure 6). Furthermore, there was a significantly negative genetic correlation between the age of onset GWAS summary statistics and the most recent PD GWAS (Chang et al., 2017) using LDSC (rg=−0.5511, se=0.103, p=9.027E-8). When solely looking at the 44 SNPs that were genome-wide significant in the most recent PD GWAS (Chang et al., 2017), we identified a significant effect in six loci after Bonferroni correction: *SNCA*, *TMEM175*, *BST1*, *INPP5F/BAG3*, *FAM47E/SCARB2*, and *MCCC1* (Figure 4, Supplemental Table 3 and Supplementary Figure 12). Interestingly, for some well-established loci no significant P-values were identified in any of the datasets, including *GCH1* (P_meta= 0.9769, 80% power to detect a change in AOO >0.28 years), *RAB7L1*/NUCKS1 (PARK16) (P_meta=0.124, 80% power to detect a change in AOO >0.26 years), and *MAPT* (P_meta=0.745, 80% power to detect a change in AOO >0.32 years) (Figure 5) (see Supplementary Table 3 for all power calculations).

**Figure 6:**
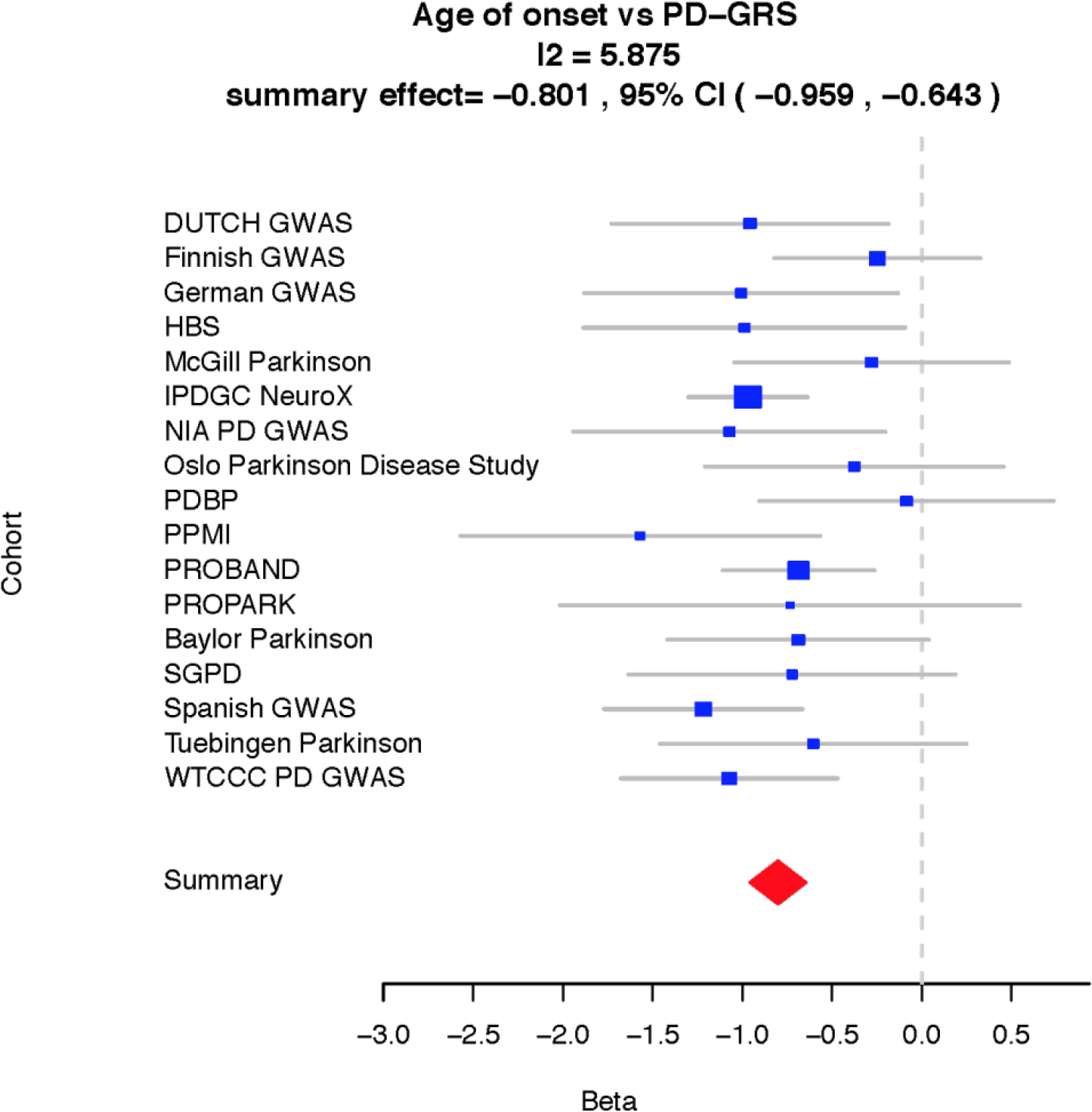
Forest plot of genetic risk score of the Parkinson’s disease GWAS correlated with the Parkinson’s disease age of onset per dataset. Overall, a consistent association is seen between the genetic risk score and age of onset. IPDGC = International Parkinson Disease Genomics Consortium, NIA = National institute on aging, HBS = Harvard Biomarker Study, PDBP = Parkinson’s Disease Biomarkers Program, PPMI = Parkinson’s Progression Markers Initiative, SGPD = System Genomics of Parkinson’s Disease

### APOE E4 allele as positive control longevity marker

Interestingly, the variant representing the *APOE* E4 allele (chr19:45411941, rs429358) was among the borderline significant loci with a P_meta value of 5.696E-8 resulting in a 0.7 year earlier of age of onset (beta=−0.707, SE=0.130). The *APOE* E4 allele is the most common genetic risk factor for Alzheimer’s disease (Liu et al., 2013), but has no effect on PD (P-value in latest GWAS=0.625) (Federoff et al., 2012; Nalls et al., 2014). Interestingly, it has already been previously identified as an aging marker (Broer et al., 2015; Garatachea et al., 2014). To investigate this in our data we performed a GWAS on the reported age of controls from the IPDGC datasets and this demonstrated a result consistent with that seen in the PD cases: P value of 1.49E-5 (Effect=−0.644, SE=0.149), suggesting that associations seen at this locus are general age related effects (Figure 5). Conditional tests based on rs429358 showed that the whole signal is coming from the *APOE* E4 allele (Supplementary Figure 10). Additionally, none of the PD loci tested has a significant P value after correction for multiple testing (Supplemental Table 5).

### Replication of previous associated loci and potential sex effects

In the past decade, several studies have performed PD age of onset analyses and nominated many genes and variants of interest (Supplemental Table 2). Many of these variants and genes were identified in smaller datasets and were reported with nominal P-values or could not be replicated in other datasets. In our current GWAS we replicate the findings in *SNCA*, *GBA* and *TMEM175* as described above (Supplemental Table 2).

Additionally, some evidence exists there that there might be sex differences in age of onset. To investigate this, we divided the IPDGC datasets (excluding the SGPD dataset) into a males-only dataset (N=11,411) and a females-only dataset (N=6,621) and performed sex-specific age of onset GWASes and meta-analyses. No genome wide-significant hits were identified in either the male or the female meta-analysis and the effect sizes for the most recent PD GWAS variants Chang et al were similar (ρ=0.532, P=0.00034, Supplementary Table 4). Interestingly, we did identify a similar sex-specific effect of the *COMT* coding variant rs4680 (p.V158M) as reported previously: P=0.0133 in males (Effect=0.370, SE=0.150) and P=0.922 in females (Effect=−0.0188, SE=0.193) (Klebe et al., 2013).

## Discussion

Here we performed a genome-wide age of onset analysis using 28,568 PD cases and discovered several PD loci associated with age of onset. We identified two genome-wide significantly associated loci—*SNCA* (P_meta = 1.90E-12) and *TMEM175* (P_meta = 9.62E-9)—as well as several sub-significant loci including *GBA*, *SCARB2*, and *BAG3*. Notably, this is the first study reporting genome-wide significance for PD GWAS loci for PD age of onset. The heritability estimate of PD age of onset was ~0.11, which is much less than the heritability of PD case-control status ~0.27 (Keller et al., 2012). Our heritability estimate was lower than an estimate from a previous study in familial PD, although this was based on a relatively small dataset (N=504 families) (Hamza and Payami, 2010). This is likely, in large part, due to the subjective nature of this phenotype.

It has been shown that PD case-control GWAS loci are associated with PD age of onset (Nalls et al., 2015). We found that a GRS based on the latest PD GWAS (Chang et al., 2017) was also highly associated with an earlier age of PD onset (Figure 6). Furthermore, there was a significant negative genetic correlation between the PD case-control GWAS and the PD age of onset GWAS. Six individual loci remain significant after Bonferroni correction, including two well-established PD loci—SNCA and *TMEM175—*and four less studied PD loci—*BST1*, *INPP5F/BAG3*, *FAM47E/SCARB2*, and *MCCC1*. Interestingly, no effect was identified for several well-established loci including *GCH1*, *RAB7L1* (PARK16) and *MAPT* (Supplemental Table 3). For these three loci, our study had 80% power (at P=0.05) to detect changes greater than 0.28, 0.26, and 0.32 years, respectively. For comparison, the two genome-wide significant hits in *SNCA* and *TMEM175* modified age of onset by 0.57 and 0.61 years. We believe this provides compelling evidence that only a subset of PD case-control risk SNPs also modulates PD age of onset, and notably that variability at *MAPT*, a major GWA identified risk factor for PD does not influence age at onset. We believe these data may suggest that different PD loci modulate risk via different pathways and that this likely has therapeutic consequences.

Interestingly, three of our top hits (*SNCA*, *TMEM175*, *and GBA*) are strongly implicated in α-synuclein mechanisms of PD pathogenesis pathology. *SNCA* encodes for the α-synuclein protein which is the major constituent of Lewy bodies, the defining pathology of PD. Both common and rare variants at *SNCA* are associated with increased PD risk, including duplications or triplications of the genomic locus (Singleton et al., 2003). One of the distinguishing features of autosomal dominant PD caused by *SNCA* gene multiplications is an early age of onset (Konno et al., 2016; Ross et al., 2008). The common variants at *SNCA* associated with PD risk and age of onset have also been demonstrated to enhance *SNCA* gene expression (Soldner et al., 2016). Thus, based on the genetics of both Mendelian and sporadic PD *SNCA* dosage appears to be a major driver of disease onset age. *TMEM175* has been shown to impair lysosomal and mitochondrial function and increases α-synuclein aggregation (Jinn et al., 2017). *GBA* has been shown to be involved in several α-synuclein aggregation mechanisms, including autophagy and enhancing α-synuclein cell to cell transmission (Bae et al., 2014; Du et al., 2015; Mazzulli et al., 2011). Other loci identified by our PD age of onset analyses have also been linked to alpha-synuclein clearance, for example for *SCARB2*, encoding the ER-to-lysosome transporter of *GBA*, and *BAG3* (Cao et al., 2017; Rothaug et al., 2014). Taken together, it is tempting to speculate about the direct link between PD age of onset loci influencing the α-synuclein accumulation and clearance pathways and the other non-significant PD loci acting in another pathway by a different mechanism, but more experiments are needed to confirm this.

Several previous studies have performed PD age of onset analyses in smaller datasets. We were unable to replicate the majority of these previous reported variants, likely because of the lack of power to detect a true signal in the smaller studies (<2,000 samples) (see Supplementary Table 2). We may have been unable to replicate previous work by Hill-Burns et al. since much of their analysis was restricted to familial PD cases (Hill-Burns et al., 2016). Their analysis of sporadic PD included almost 4,000 cases, but unlike our study and several others (Brockmann et al., 2013; Lill et al., 2015; Nalls et al., 2015) they did not find significant associations with *SNCA* (rs356203, P=0.355) or *TMEM175* (rs34311866, P=0.525).

Although we included a very large amount of data there were still several limitations to our study. First, we had limited power to detect rare variants that modulate PD age of onset due to the large number of cohorts with a relatively small size. Several reports have shown that rare variants may influence age of onset of PD. For example, a recent report showed that rare variants in *LRRK2* lowered age of onset (Xiao et al., 2018). This likely also explains the relatively moderate P-values for rare coding variants in *GBA* and *LRRK2* on age onset. Second, due to the lack of reliable genotype data we excluded chromosomes X and Y from our analysis. Both of these limitations could be addressed by using whole-genome sequencing data and large whole-genome sequencing projects are currently underway. Third, the heritability of PD age of onset (~0.11) was much lower than the heritability of PD case-control status (Keller et al., 2012), which in part explains the lack of genome-wide significantly associated loci in our study. Fourth, there was a reasonable amount of heterogeneity in the mean age of onset our different cohorts. Age of PD diagnosis was self-reported in some of our cohorts and assessed by physicians in other cohorts. Some cohorts were specifically designed to include younger onset cases. Other cohorts provided age of diagnosis while others used age of onset, although these are likely to be highly correlated (r>0.9 in our data). By analyzing each cohort separately, we were able to mitigate many of these inter-study sources of variation. Indeed, there was relatively little heterogeneity of SNP effect size estimates between studies (Supplementary Table 3). However, future data collection and studies would probably benefit from more specific and more predefined structured and symptom specific age of onset diagnostic criteria. Implementation of such criteria in large studies and cohorts or even in healthcare systems could significantly improve the understanding of the genetics of PD age of onset.

Overall, we have performed the largest age of onset of PD GWAS to date and our results reveal an interesting and significant genetic component. Our results show that not all PD risk loci influence age of onset with significant differences between risk alleles for age at onset, which implies different mechanisms for risk for developing PD and PD age of onset. This provides a compelling picture, both within the context of functional characterization of disease linked genetic variability and in defining differences between risk alleles for age at onset, or frank risk for disease. The significantly associated variability is centered on the gene encoding alpha-synuclein and on variability in several other lysosomal proteins that have been shown to directly influence alpha-synuclein aggregation or clearance. Thus, these data support the notion that alpha-synuclein is an important target for future disease modifying or preventive therapies and that drugs targeting alpha-synuclein production and clearance may be most valuable.

## Acknowledgements

We would like to thank all of the subjects who donated their time and biological samples to be a part of this study. We also would like to thank all members of the International Parkinson Disease Genomics Consortium (IPDGC). See for a complete overview of members, acknowledgements and funding http://pdgenetics.org/partners. We also thank the research participants and employees of 23andMe for making this work possible. This work was supported in part by the Intramural Research Programs of the National Institute of Neurological Disorders and Stroke (NINDS), the National Institute on Aging (NIA), and the National Institute of Environmental Health Sciences both part of the National Institutes of Health, Department of Health and Human Services; project numbers 1ZIA-NS003154, Z01-AG000949-02 and Z01-ES101986. In addition, this work was supported by the Department of Defense (award W81XWH-09-2-0128), and The Michael J Fox Foundation for Parkinson’s Research. This work was supported by National Institutes of Health grants R01NS037167, R01CA141668, P50NS071674, American Parkinson Disease Association (APDA); Barnes Jewish Hospital Foundation; Greater St Louis Chapter of the APDA. The KORA (Cooperative Research in the Region of Augsburg) research platform was started and financed by the Forschungszentrum für Umwelt und Gesundheit, which is funded by the German Federal Ministry of Education, Science, Research, and Technology and by the State of Bavaria. This study was also funded by the German Federal Ministry of Education and Research (BMBF) under the funding code 031A430A, the EU Joint Programme - Neurodegenerative Diseases Research (JPND) project under the aegis of JPND -www.jpnd.eu- through Germany, BMBF, funding code 01ED1406 and iMed - the Helmholtz Initiative on Personalized Medicine. This study is funded by the German National Foundation grant (DFG SH599/6-1) (grant to M.S), Michael J Fox Foundation, and MSA Coalition, USA (to M.S). The McGill study was funded by the Michael J. Fox Foundation and the Canadian Consortium on Neurodegeneration in Aging (CCNA). This study utilized the high-performance computational capabilities of the Biowulf Linux cluster at the National Institutes of Health, Bethesda, MD, USA. (http://biowulf.nih.gov), and DNA panels, samples, and clinical data from the National Institute of Neurological Disorders and Stroke Human Genetics Resource Center DNA and Cell Line Repository. People who contributed samples are acknowledged in descriptions of every panel on the repository website. We thank P Tienari (Molecular Neurology Programme, Biomedicum, University of Helsinki), T Peuralinna (Department of Neurology, Helsinki University Central Hospital), L Myllykangas (Folkhalsan Institute of Genetics and Department of Pathology, University of Helsinki), and R Sulkava (Department of Public Health and General Practice Division of Geriatrics, University of Eastern Finland) for the Finnish controls (Vantaa85+ GWAS data). We used genome-wide association data generated by the Wellcome Trust Case-Control Consortium 2 (WTCCC2) from UK patients with Parkinson’s disease and UK control individuals from the 1958 Birth Cohort and National Blood Service. Genotyping of UK replication cases on ImmunoChip was part of the WTCCC2 project, which was funded by the Wellcome Trust (083948/Z/07/Z). UK population control data was made available through WTCCC1. This study was supported by the Medical Research Council and Wellcome Trust disease centre (grant WT089698/Z/09/Z to NW, JHa, and ASc). As with previous IPDGC efforts, this study makes use of data generated by the Wellcome Trust Case-Control Consortium. A full list of the investigators who contributed to the generation of the data is available from www.wtccc.org.uk. Funding for the project was provided by the Wellcome Trust under award 076113, 085475 and 090355. This study was also supported by Parkinson’s UK (grants 8047 and J-0804) and the Medical Research Council (G0700943 and G1100643). Sequencing and genotyping done in McGill University was supported by grants from the Michael J. Fox Foundation, the Canadian Consortium on Neurodegeneration in Aging (CCNA) and in part thanks to funding from the Canada First Research Excellence Fund (CFREF), awarded to McGill University for the Healthy Brains for Healthy Lives (HBHL) program. PRoBaND data was funded by Parkinson’s UK (Grant ref J-1101) and supported by NHS Greater Glasgow & Clyde and University of Glasgow. DNA extraction work that was done in the UK was undertaken at University College London Hospitals, University College London, who received a proportion of funding from the Department of Health’s National Institute for Health Research Biomedical Research Centres funding. This study was supported in part by the Wellcome Trust/Medical Research Council Joint Call in Neurodegeneration award (WT089698) to the Parkinson’s Disease Consortium (UKPDC), whose members are from the UCL Institute of Neurology, University of Sheffield, and the Medical Research Council Protein Phosphorylation Unit at the University of Dundee. We thank the Quebec Parkinson’s Network (http://rpq-qpn.org) and its members. This work was supported by the Medical Research Council grant MR/N026004/1. Data used in the preparation of this article were obtained from the Parkinson’s Progression Markers Initiative (PPMI) database (www.ppmi-info.org/data). For up-to-date information on the study, visit www.ppmi-info.org.

PPMI, a public-private partnership, is funded by the Michael J. Fox Foundation for Parkinson’s Research and funding partners, including AbbVie, Avid, Biogen, Bristol-Myers Squibb, Covance, GE Healthcare, Genentech, GlaxoSmithKline, Lilly, Lundbeck, Merck, Meso Scale Discovery, Pfizer, Piramal, Roche, Servier, Teva, UCB, and Golub Capital. Data and biospecimens used in preparation of this manuscript were obtained from the Parkinson’s Disease Biomarkers Program (PDBP) Consortium, part of the National Institute of Neurological Disorders and Stroke at the National Institutes of Health. Investigators include: Roger Albin, Roy Alcalay, Alberto Ascherio, DuBois Bowman, Alice Chen-Plotkin, Ted Dawson, Richard Dewey, Dwight German, Xuemei Huang, Rachel Saunders-Pullman, Liana Rosenthal, Clemens Scherzer, David Vaillancourt, Vladislav Petyuk, Andy West and Jing Zhang. The PDBP Investigators have not participated in reviewing the data analysis or content of the manuscript. A full acknowledgement is available in the Supplementary data.

## Author Contributions

### Data Acquisition or data contribution

CB, KH, CLV, SBC, RC, LP, JSS, CS, MS, LK, AS, HI, HL, AJN, JRG, DGH, SWS, JJ, LMS, SL, JCC, AB, JJH, JM, PT, KM, MT, DGG, TG, PH, JMS, NW, JH, HRM, DAH, JG, PMV, ZGO, MAN, ABS

### Study level analysis and data management

CB, KH, CLV, MAN

### Design and funding

TG, PH, JMS, NW, JH, HRM, JG, PMV, ZGO, MAN, ABS

### Writing – Original Draft

CB, KH, MAN, ABS

### Critical review and writing the manuscript

CB, KH, CLV, SBC, RC, LP, JSS, CS, MS, LK, AS, HI, HL, AJN, JRG, DGH, SWS, JJ, LMS, SL, JCC, AB, JJH, JM, PT, KM, MT, DGG, TG, PH, JMS, NW, JH, HRM, DAH, FG, PMV, ZGO, MAN, ABS

